# Flexible Control as Surrogate Reward or Dynamic Reward Maximization

**DOI:** 10.1101/2022.02.25.482039

**Authors:** Mimi Liljeholm

## Abstract

The utility of a given experience, like interacting with a particular friend or tasting a particular food, fluctuates continually according to homeostatic and hedonic principles. Consequently, to maximize reward, an individual must be able to escape or attain outcomes as preferences change, by switching between actions. Recent work on human and artificial intelligence has defined such flexible instrumental control in information theoretic terms and postulated that it may serve as a reward surrogate. Another possibility, however, is that the adaptability afforded by flexible control is tacitly implemented by planning for dynamic changes in outcome values. In the current study, an expected utility model that computes decision values over a range of possible monetary gains and losses associated with sensory outcomes provided the best fit to behavioral choice data and performed best in terms of earned rewards. Moreover, consistent with previous work on perceived control and personality, individual differences in dimensional schizotypy were correlated with behavioral choice preferences in conditions with the greatest and lowest levels of flexible control. These results contribute to a growing literature on the role of instrumental control in goal-directed choice.

## 1. Introduction

What does it mean to control your environment and why is it important? A substantial body of research has demonstrated a preference for voluntary choice, across a wide range of animal species, from pigeons and rats to monkeys and humans (1; 2; 3; 4; 5). Specifically, even when the values and probabilities of outcomes are equated, subjects prefer environments in which they are free to choose between action alternatives over ones in which they are forced to select a particular action. Such findings pose a problem for normative theories of economic choice, which postulate that preferences are determined solely by the probabilities and subjective utilities of decision outcomes. A rational explanation for the free-choice preference is that subjective outcome utilities often change from one moment to the next, and that free choice allows an agent to maximize long run rewards by switching between actions to flexibly produce whichever outcome is most preferred at any given time.

### Box 1

**Glossary**

**Subjective Utility:** The perceived satisfaction conferred by a commodity or service.

**Expected Utility:** An expectation of utility based on the conditional probability and subjective utility of outcome states.

**Instrumental Divergence:** The degree to which freely chosen action alternatives yield distinct sensory consequences.

**Information Theoretic (IT) Distance:** A quantitative specification of the dissimilarity between probability distributions.

**Outcome predictability:** The uniformity of the outcome probability distribution (i.e., entropy).

**Outcome diversity:** The number of distinct and obtainable outcomes.

**Sense of Agency (SOA):** The subjective experience of controlling one’s actions and their consequences.

**Positive Schizotypy:** A multidimensional personality trait characterized by unusual perceptual experiences and odd beliefs.

However, recently, it has been noted that free choice between actions that produce highly similar or identical outcomes affords no such flexibility and that, consequently, *instrumental divergence –* the degree to which freely chosen actions yield distinct consequences – is an essential aspect of dynamic reward maximization (6; 7; 8; 9). Formally, the *information theoretic* (IT) distance between outcome probability distributions associated with action alternatives has been proposed to capture an agents control over its environment (6; 7; 8; 9; 10). Of course, without free choice, the IT distance between outcome distributions, although related to the predictability and diversity of outcomes, would not be instrumental and, consequently, would have no implications for flexible instrumental control. Greater instrumental divergence, thus, reflects *both* free choice and greater IT distance between outcome distributions (6; 7; 8; 9). Moreover, IT variables do not necessarily capture the full scope of instrumental flexibility – indeed, an explicit representation of instrumental divergence, however formalized, may not be necessary for adaptive choice. Here, a more tacit role of instrumental divergence is probed, as an estimation of decision values that adjusts for dynamic outcome utilities.

As an illustration, imagine that you are a gambler choosing which of two “rooms” to gamble in, knowing that you will only have access to the slot machines in the selected room for several rounds of gambling. Imagine, further, that each slot machine yields three differently colored outcomes with various probabilities, and that the monetary amounts associated with the different outcomes, which can be either positive or negative, are currently unknown. Now, consider the two room options illustrated in the top panel of Figure 1 (ignore self- vs. auto-play labels for the purpose of this example): In the Left Room, the two available slot machines, depicted as pie charts, have identical distributions across the three outcome colors. Accordingly, the IT distance between the outcome probability distributions of the two machines is zero. In the Right Room, one of the machines uniquely delivers red tokens, yielding a slightly greater, but still low, IT distance between token probability distributions. Critically, while the distance between outcome distributions associated with available gambling options is low in both rooms, the Right Room uniquely affords complete avoidance of the red outcome color, which is useful whenever the red token is associated with a negative value: Conversely, in the bottom panel of Figure 1, both rooms have relatively large distances between outcome distributions associated with alternative gambling options, again yielding a small difference between rooms, and both rooms afford the opportunity to completely avoid a particular outcome color (i.e., red).

**Figure 1:**
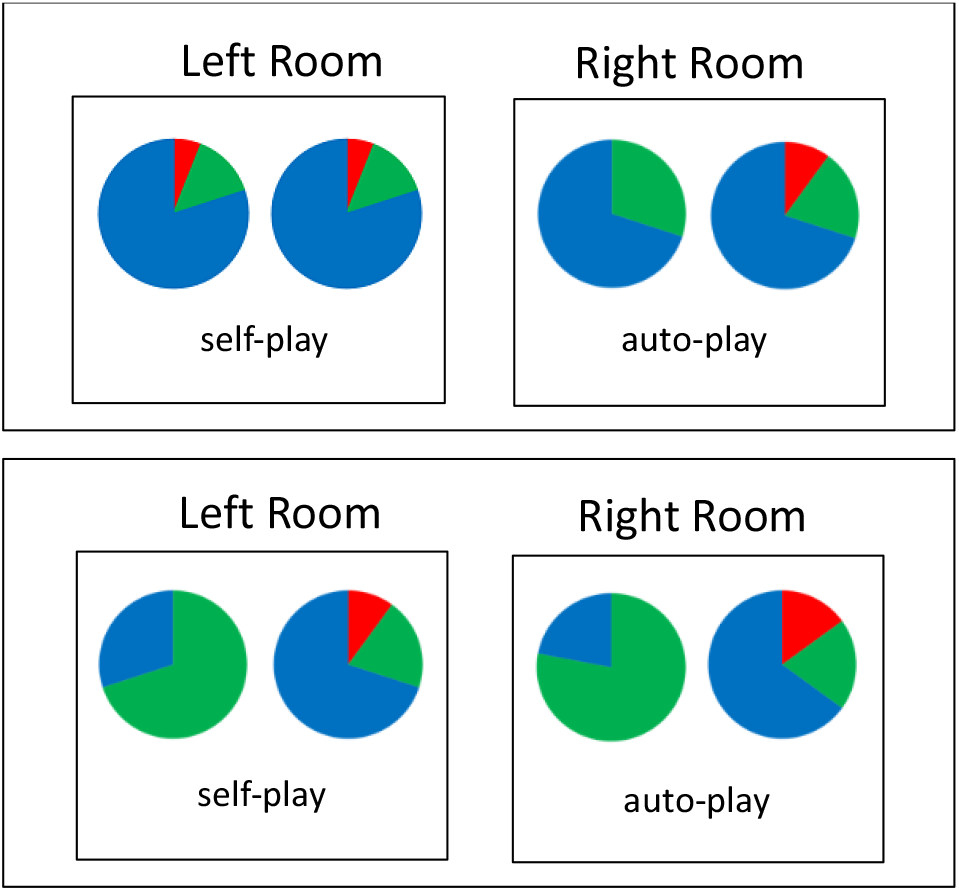
Task illustration. Choice screen shown at the beginning of a gambling round, where “room” options differ in terms of the information theoretic (IT) distance between color outcome distributions (represented by pie charts) associated with the “slot machines” in each room, and in terms of whether gambling in the room involves free (self) or forced (auto) choice.

In other words, based on the IT distance between token probability distributions, there should be no difference in the Right-over-Left Room preference across top and bottom panels in Figure 1. However, if representations of instrumental divergence involve a consideration of the avoidability of potentially aversive outcomes, the preference for the Right Room should be greater in the top panel. Here, to assess their normative and descriptive performance, both approaches – using IT distance as a reward surrogate vs. planning for potentially negative outcome utilities – are instantiated in an expected utility framework.

Importantly, while instrumental divergence is an affordance of the environment, it is filtered through the mind of an agent whose beliefs, discernments, and biases play a critical role in decision making. Substantial evidence suggests that positive symptoms of schizophrenia (e.g., hallucinations and delusions) are associated with an exaggerated *sense of agency* (SoA) – defined by (11) as “the experience of controlling one’s own motor acts and, through them, the course of external events” – whether assessed using declarative statements of self- vs. external attribution (12; 13; 14), or implicitly, as altered time perception associated with voluntary actions (15) or interference by another’s movements on motor performance (16). Given the correspondence between instrumental divergence and contemporary accounts of agency, see (9) for review, this suggests that positive symptoms may also dysregulate a preference for instrumental divergence: Here, individual differences in dimensional positive schizotypy were used to predict the preference for greater instrumental divergence in neurotypical adults.

## 2. Methods

### 2.1 Participants

One hundred and twenty individuals (45 females; mean age = 23.7 ± 3.6) participated on MTurk (Amazon Mechanical Turk) for a $5 baseline compensation and up to $7 in additional earnings based on experimental contingencies. A power analysis performed on an independent sample (n=50), using G*Power 3.1 (17), revealed that a sample size of 115 would be required to detect a significant difference between self- and auto-play conditions in the preference for greater IT distances with a power of 0.90. All participants gave informed consent, and the study was approved by the Institutional Review Board of the University of California, Irvine.

### 2.2 Task & Procedure

The task is illustrated in Figures 1 & 2. At the start of the experiment, participants were instructed that they would assume the role of a gambler playing various slot machines in a casino, with the goal of maximizing the amount of monetary gain. Participants were further told that, in each of several gambling rounds, they would be required to first select a room in which only two slot machines were available, that they would be restricted to gamble only on the machines available in the selected room on several trials within that round, and that they would get to keep the monetary earnings, up to $7, from three gambling rounds, randomly drawn from all rounds at the end of the study.

**Figure 2:**
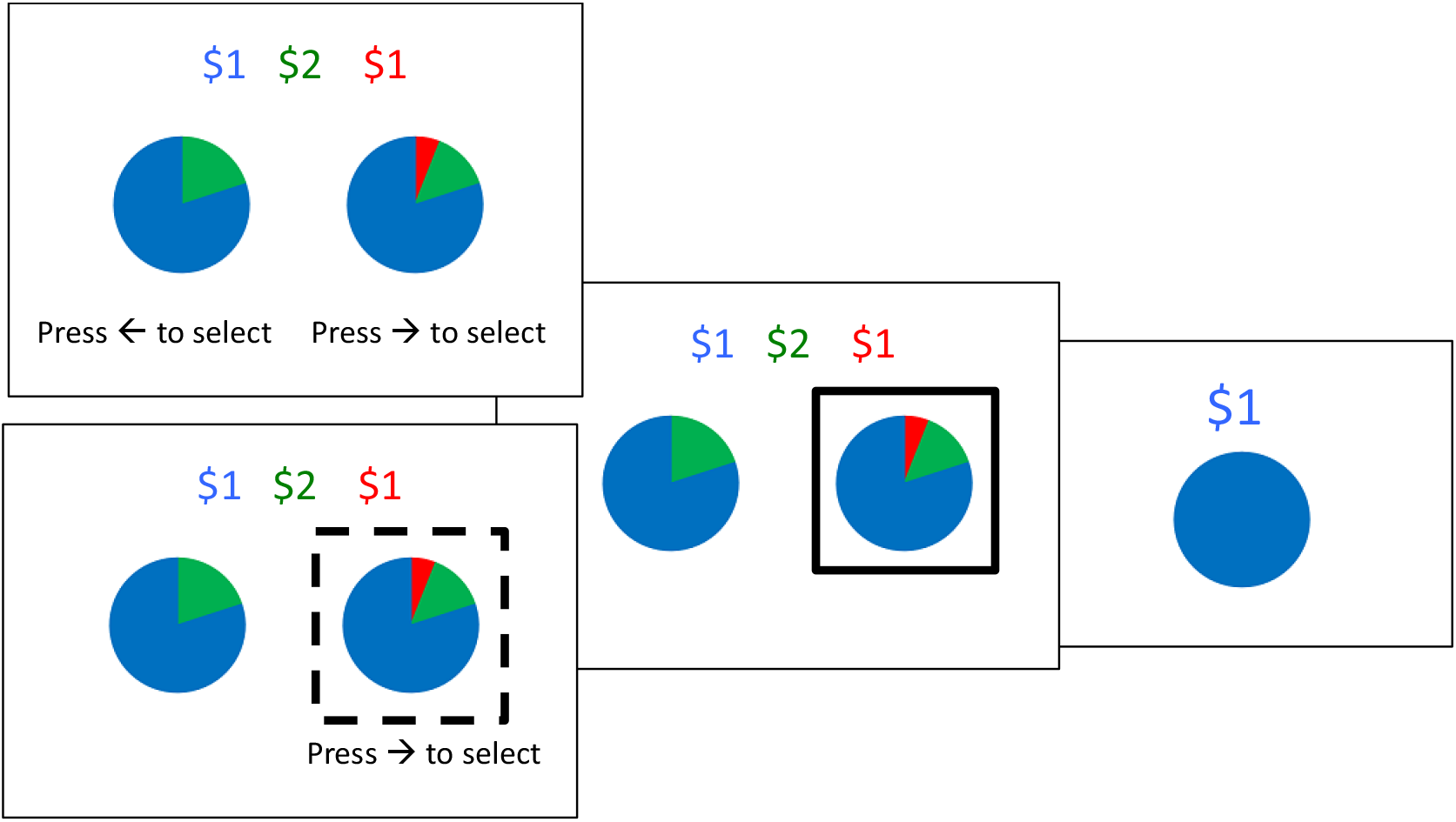
Task illustration. Choice, selection & feedback screens on a trial inside a gambling room. In auto-play conditions, a dashed square around one gambling option indicates the forced choice (bottom left). Once a key is pressed, chosen freely (self-play) or as instructed by the dashed frame (auto-play), a solid-line square frames the selection for 1 second, followed by a 2-second display of the obtained outcome. Note that, once inside the room, the current values of the outcome colors are displayed at the top of the choice screen. These values stay the same throughout the gambling round but change once the participant selects a new room to gamble in (see choice screens in Figure 1).

Each slot machine yielded three different colors (red, green, and blue) with different probabilities, and each color was worth a particular monetary amount, which changed from gambling round to gambling round, and which was only revealed once a room had been selected. The probabilities with which a given slot machine produced the red, green, and blue outcome were graphically illustrated using pie chart slices. The primary measure was the decision at the beginning of each round (see Figure 1), between gambling rooms that differed in terms of the IT distance between outcome distributions associated with available slot machines.

Recall that instrumental divergence refers to the difference between the consequences of *freely chosen* action alternatives. Here, to experimentally dissociate the role of flexible control from that of outcome diversity (i.e., the variability of obtainable outcomes) or predictability, we use a self- vs. auto-play manipulation, such that all room options were either self-play – participants choose freely between slot machines available in the selected room – or auto-play – a computer algorithm alternated between machines across trials in the selected room – as indicated by labels printed below options on the room-choice screen in Figure 1. Whereas the difference between rooms in terms of outcome diversity and predictability is the same for self- and auto-play rooms, instrumental divergence is always zero in auto-play rooms. Importantly, the self- vs. auto-play manipulation also allows us to assess whether the frequently demonstrated preference for free-choice (6; 7; 8; 9) is modulated by the flexibility of control (i.e., instrumental divergence).

The difference between outcome distributions associated with the slot machines in a room was formalized as the Information Theoretic (IT) distance (see glossary and section 2.3) of the room, with 4 such distances (0.00, 0.04, 0.15 and 0.20) yielding 6 unique “IT distance-differences” – that is, differences *between rooms* in the IT distance between outcome distributions associated with the slot machines in each room; specifically, 0.00 vs 0.04, 0.00 vs 0.15, 0.00 vs 0.20, 0.04 vs 0.15, 0.04 vs 0.20, and 0.15 vs 0.20. IT-distance conditions were combined with 4 self- vs auto-play combinations (both rooms self-play, both rooms auto-play, greater divergence room self-play, lesser divergence room self-play), which were repeated across two color schemes, for a total of 48 room choice trials.

Once a room had been selected, there were 4 discrete slot machine choices performed in that room, yielding a total of 256 gambling trials. The monetary values of color outcomes differed across rounds of gambling, and participants were explicitly instructed that these values would “change from round to round”. To ensure that stochastically generated monetary payoffs could not account for a preference for a greater IT distance, reward distributions were constructed such that monetary pay-offs were largely balanced, and, if anything, biased *against* the hypothesized influence of the IT distance. The programmed expected monetary payoffs, with the expectation taken over the products of the probability and $ amount associated with each outcome color for each slot-machine, (using the max for self-play rooms and mean for auto-play rooms) are listed for each IT distance and self- v. auto-play condition in Table 1.

**Table 1:** Mean Shannon entropy and expected monetary payoffs, given self- and auto-play respectively, at each divergence level.

| Room Divergence | 0.00 | 0.04 | 0.15 | 0.20 |
| --- | --- | --- | --- | --- |
| Mean Entropy | 0.74 | 0.71 | 0.71 | 0.71 |
| Self-play pay-off | \$0.39 | \$0.36 | \$0.34 | \$0.38 |
| Auto-play pay-off | \$0.39 | \$0.37 | \$0.34 | \$0.37 |

### 2.3 Computational Variables

Under some conditions, instrumental divergence may be defined as the IT distance (19) between the outcome probability distributions associated with available actions. Let *P*_*1*_ and *P*_*2*_ be the respective color probability distributions of the two slot machines available in a given room, let *O* be the set of possible color outcomes, and *P(o)* the probability of a particular color outcome. The IT distance is:

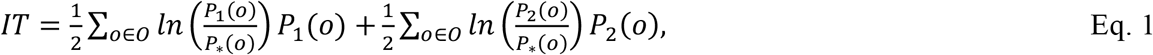

where ln is the natural logarithm, yielding nats, and

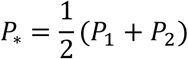

Note that instrumental divergence pertains to the sensory, rather than motivational, features of outcome states – since subjective utilities are constantly changing, motivational features are intrinsically unstable as a basis for estimating flexible control – and with respect to outcome distributions associated with *available* action alternatives, rather than observed actions or cues – since only freely chosen actions confer flexible instrumental control. Thus, while an IT distance can be computed over probability distributions associated with any type of random variable, it only indicates *instrumental* divergence when it is computed over sensory outcome probability distributions associated with freely chosen actions.

Three expected utility models were tested that instantiate different normative and descriptive hypotheses regarding the quantitative integration of the utility of instrumental divergence with conventional monetary reward. First, in all three models, the acquired value of a particular gambling room (i.e., a particular pair of pie charts) was incrementally updated across gambling rounds, such that

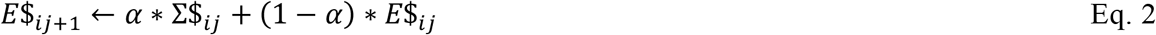

where *E*$_*ij*+1_ is the expected monetary pay-off in room *i* at the start of round *j+1* (i.e., at the start of the next round played in room *i*), ∑$_*ij*_ is the sum of monetary outcomes earned in room *i* at the end of round *j* (i.e., the end of the current round), *E*$_*ij*_ is the expected monetary payoff in room *i* on round *j* (recursively estimated based on the payoff, ∑$_*ij*−1_, and expectation, *E*$_*ij*−1_, in the previous round, *j-1*), and *a* is a free learning rate parameter. In other words, despite explicit instructions that the monetary values of different token colors will change from round to round, all models assume that expectations about monetary outcomes depend, to some extent, on experienced payoffs. In model simulations, ∑$_*ij*_ was generated using the monetary values and probabilities of outcomes in each chosen room, assuming optimal selection of slot-machines in self-play rooms and alternation across slot-machines in auto-play rooms.

All three models also included a free parameter, *γ*, quantifying the subjective utility of free over forced choice (i.e., self-play). The models differed, however, in their treatment of instrumental divergence. In the first model, the value of a particular room, *V_SP_*, informing the decision of which room to gamble in at the onset of each trial, depended only on the incrementally updated expected monetary payoff, and whether the room was self- or auto-play:

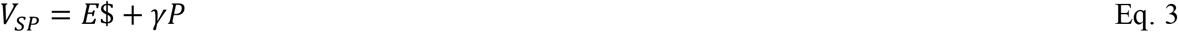

where *P* is an indicator set to 1 for self-play and 0 for auto-play. In a second model, *V_IT_*, the decision value of a room depended additionally on the subjective utility of the information theoretic distance, *D*, between token probability distributions associated with the two slot machines (i.e., pie charts) available in a room, such that

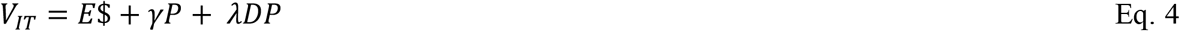

where λ is a free parameter. Note that V_IT_ scales with the IT distance in self-play rooms only; in auto-play rooms, in which IT distance does not correspond to instrumental divergence, there is no influence of IT distance on V_IT_ (i.e., *P* equals 0).

Finally, in a third model, V_FWD_, the value of instrumental divergence is captured, not in terms of an IT distance, but by simulating a range of possible token values and computing a forward estimate of the room payoff:

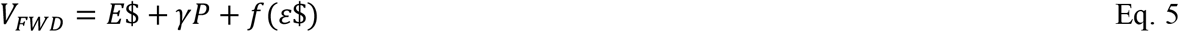

where *ε*$ averages across the products of the probabilities indicated by the pie charts and the *possible* values of the different token colors, such that

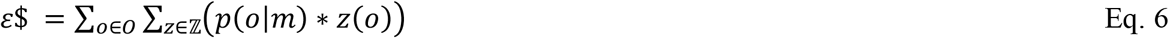

where *p(o*|*m)* is the probability of a particular token outcome, *o*, given selection of a particular slot machine, *m*, and *z(o)* is a possible value of token *o* from the set ℤ = {−$3: $1: $3}. Critically, when a room is self-play (i.e., P=1), the slot machine with the greatest payoff given the combination of values from ℤ is used to compute *ε*$; conversely, when a room is auto-play, computations over possible token values are averaged across machines, thus always summing to zero, reflecting the alternating selection of machines across trials in the room:

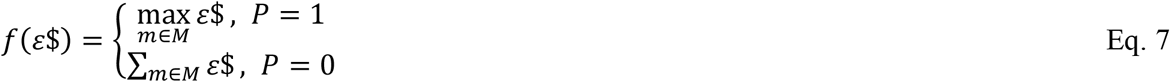

In other words, rather than being explicitly represented (e.g., as an IT distance) and treated as a reward surrogate, the forward model maximizes reward across possible outcome values, given the IT distance, the presence vs. absence of free choice, and the obtainability and avoidability of outcomes.

Model-derived room values (*V*_*room*_) were transformed into room choice probabilities using a softmax rule with a noise parameter, τ;

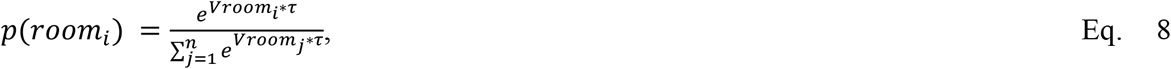

The Shannon entropy of a slot machine available in a room (i.e., the “flatness” of its outcome probability distribution) was defined as:

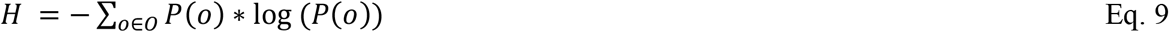

Free model parameters were fit to behavioral data by minimizing the negative log likelihood and computing the corrected Akaike Information Criterion (AIC). All computational variables were implemented using MATLAB (https://www.mathworks.com/).

### 2.4 Assessment of Positive Schizotypy

The Oxford-Liverpool Inventory of Feelings and Experiences (O-LIFE) is a four-scale questionnaire intended to assess dimensions of schizotypy, with scales corresponding, respectively to “unusual experiences”, “cognitive disorganization”, “introvertive anhedonia”, and “impulsive non-conformity”. In particular, the “unusual experiences” dimension has been phenomenologically related to positive symptoms of schizotypy (17; 18) and, thus, was of primary interest here. We hypothesized that differences along this dimension would predict individual differences in the preference for instrumental divergence (i.e., for a combination of high-divergence and self-play). The O-LIFE questionnaire was administered immediately after the gambling task for all participants. The Pearson correlation coefficient was computed between positive schizotypy and lowest (0)- vs. greatest- (0.2) divergence conditions, for self-play vs. auto-play rooms.

### 2.5 Statistical Analyses

Paired-samples two-tailed t-tests were used to compare 1) reward acquisition and choice preferences by different expected utility models, 2) the relative fits of expected utility models to behavioral choice data (i.e., model AIC scores), 3) behavioral choice preferences for greater IT-distance rooms in self- vs auto-play conditions, and 4) choice preferences across the conditions detailed in Figure 1. Effect sizes (Cohen’s *d*) are provided for all t-tests. In addition, two-tailed Pearson’s correlation coefficients were computed between the positive schizotypy measure and the preferences for greater IT distance, in self- vs. auto-play conditions. All statistical analyses were implemented in JASP (https://jasp-stats.org/).

## 3. Results

To evaluate reward acquisition and parameter recovery, a data set was simulated consisting of 1000 gamblers with 1000 room decisions each, and with color outcome values of $-3 to $3, incrementing by $1, drawn from a uniform distribution for each round. The mean monetary payoff earned by gamblers simulated with the V_FWD_ model was significantly greater than that of subjects simulated withe V_IT_ model, t(999)=4.55, *p<0*.*001, d=0*.*14*, which in turn was significantly greater than that of subjects simulated with the V_SP_ model, t(999)=7.60, *p<0*.*001, d=0*.*24*. In other words, among the models evaluated, the V_FWD_ model appears to be optimal. Across models, parameter recovery was highly significant for all parameters (p<0.0001), with excellent recovery for the learning rate parameter (r>=0.88) and robust recovery for noise (r>=0.54), self- vs. auto-play (r>=0.62), and IT-distance (r=0.48) parameters.

The model-derived probability of choosing a room is plotted, for each expected utility model, as a function of IT distance, for self- and auto-play rooms, together with mean observed choice proportions, in Figure 3. Note, first, that all models categorically discount the auto-play condition. Likewise, all models predict an influence of a room’s anticipated monetary pay-off, which varies inversely with, and thus counters the influence of, IT distance (see section 2.2 and Table 1), and which is the sole basis for *V*_*SP*_ predictions within self- and auto-play conditions. Finally, both the *V*_*IT*_ and *V*_*FWD*_ model scale with the IT distance in self- but not auto-play rooms; the former because it treats the IT-distance as a reward surrogate and the latter because greater IT-distance yields greater expected payoffs computed over the range of possible outcome utilities.

**Figure 3:**
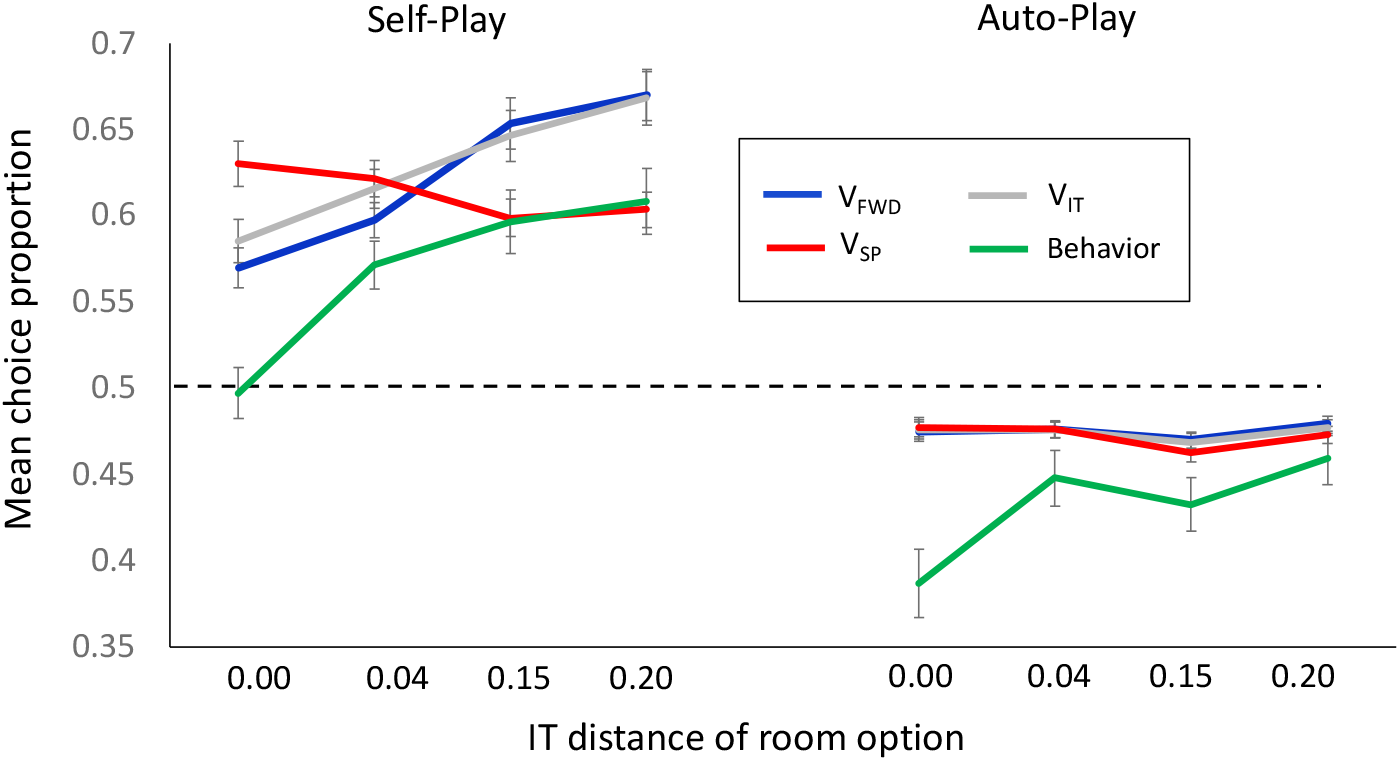
Results. Mean proportions of choosing a particular room to gamble in, as a function of the information theoretic distance between outcome distributions associated with gambling options available in the room, for behavioral choice performance and for three expected utility models. Choice proportions were averaged, for each IT-distance room, across all pairwise choice scenarios in which a room option occurred, separately for self-and auto-play rooms. All models reflect the experienced monetary payoff of a room, and the value of self- vs. auto-play (SP). The V_IT_ model also accounts for the value of greater IT distance, while the forward, V_FWD_, model considers a range of possible outcome utilities.. Dashed lines indicate chance performance and error bars = SEM.

**Figure 4:**
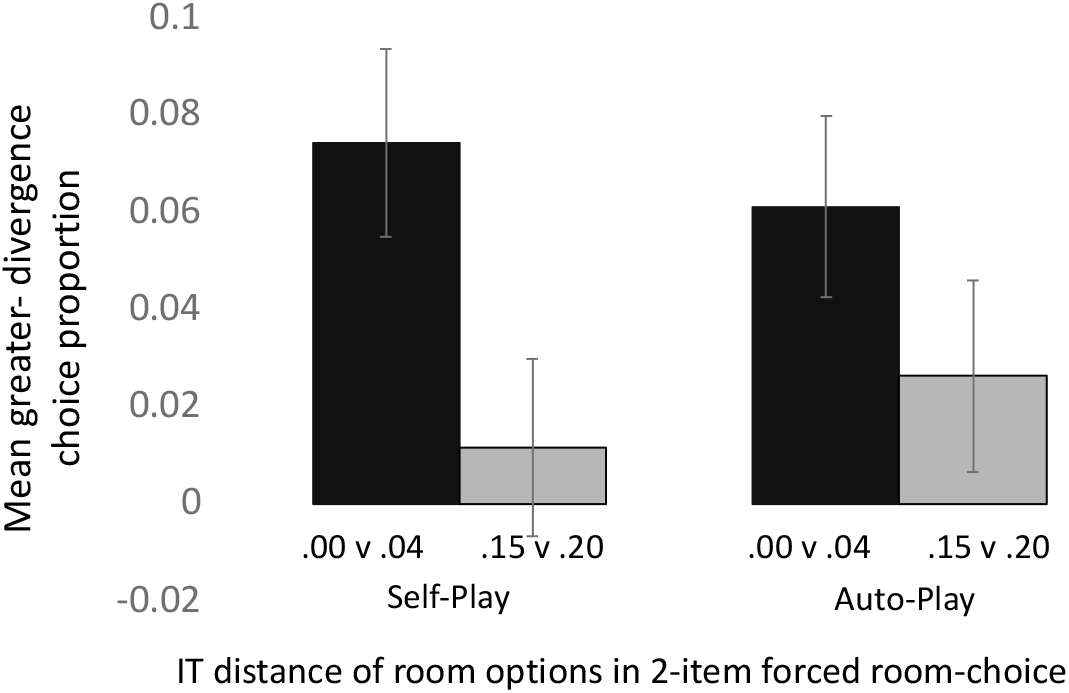
Results. Mean choice proportions across the conditions illustrated in Figure 1, showing that the increase in choice proportion with an increase in IT distance was significantly greater across rooms with low IT distance than across rooms with high IT, in the self-play but not the auto-play condition.

As can be seen in Figure 3, contrary to both the *V*_*IT*_ and *V*_*FWD*_ model, participants categorically avoided rooms with zero IT distance, whether self- or auto-play, possibly because of the slightly greater entropy in the zero-distance condition (see Table 1), or because slot machines in that condition were identical. Nevertheless, overall, the *V*_*FWD*_ model provided the best fit to behavioral choice proportions: The AIC score was significantly lower for the *V*_*FWD*_ model than for both the *V*_*IT*_(t(119)=22.65, p<0.001, *d*=2.07) and the *V*_*SP*_ (t(119)=4.49, p<0.001, *d*=0.41) model.

The probability of choosing a room with greater divergence, computed across all pairwise IT- distances, revealed that participants were more likely to select the room with the greater IT distance when both rooms were self-play (mean choice probability = 0.58) than when both rooms were auto-play (mean choice probability = 0.54; t(119)=2.17, p=0.03, d=0.20), suggesting a specific preference for instrumental control, rather than mere outcome diversity. Planned comparisons of choice proportions across the conditions illustrated in Figure 1 revealed that the increase in choice proportion with an increase in IT distance was greater across rooms with low IT distance (top panel in Figure 1) than across rooms with high IT distance (bottom panel in Figure 1); presumably because, in the former case, the increase in IT distance was associated with an opportunity to completely avoid a potentially aversive outcome. However, this differential increase in choice proportions was only significant when the room with a greater IT-distance was self-play (t(119)=2.40, p=0.018, *d=0*.*22*), not when it was auto-play, p=0.188.

Finally, given the substantial literature implicating positive symptoms of schizophrenia in an exaggerated sense of agency (12; 16; 14; 15; 13), the Short Oxford-Liverpool Inventory of Feelings and Experiences (O-LIFE; (20)) was used to assess whether positive schizotypy predicted preferences for flexible instrumental control. Across participants, positive schizotypy was *positively* correlated with the probability of choosing a zero-divergence auto-play room (r=0.28, p=0.0019) and *negatively* correlated with the probability of choosing a high-divergence self-play room (r=-0.28, p=0.0017); in contrast, schizotypy did *not* significantly predict the probability of choosing a zero-divergence self-play room (p=0.95), nor a high-divergence auto-play room (p=0.12). Finally, there was a strong negative correlation between positive schizotypy and the overall preference for self-play, collapsed across divergence conditions (r=-0.34, p<0.001).

## 4. Discussion

This study used a novel gambling task in which the decision of interest was a choice between different gambling “rooms”, given information about the divergence with which a pair of slot machines available in respective rooms produced a set of distinctly colored outcomes. Critically, in each round of gambling, the monetary values of the tokens changed, with new values being revealed only *after* a room had been selected, and with gambling restricted to the slot machines in the chosen room for several trials. In addition to outcome probability distributions associated with available slot machines, gambling rooms differed in terms of whether participants were free to choose between machines during the round (self-play) or forced to alternate between slot machines across trials (auto-play). Three expected utility models were implemented to assess whether the preference for greater instrumental divergence reflected treatment of this variable as a surrogate reward term, or a forward maximization of expected utility over *possible* future monetary payoffs. Participants’ choice behavior was best explained by the forward model, suggesting that, rather than being intrinsically rewarding, instrumental divergence is used as a planning variable.

Recent work on human (8) and artificial (10) intelligence suggests that an explicit representation of instrumental control may serve as a reward surrogate, reinforcing and motivating decisions and representations that yield high-agency states. Formally, the information theoretic (IT) distance between transition probability distributions associated with action alternatives has been proposed to capture an agents control over its environment (6). However, IT measures do not necessarily reflect an agent’s estimate of instrumental control. In particular, as illustrated in Figure 1, there are instances in which differences in IT distance do not correspond to an agent’s ability to escape (potentially aversive) outcomes. Here, a forward estimation of action values based on the range of possible outcome utilities outperformed the IT distance model with respect to both reward maximization and fit to behavioral choice data, demonstrating normative as well as descriptive advantages.

Choice preferences at different IT-distances, in self- vs. auto-play conditions, were differentially predicted by individual differences in dimensional schizotypy. Specifically, the greater an individual’s positive schizotypy score, the *more* likely that individual was to select the zero-distance auto-play room, and the *less* likely that individual was to choose the greatest-distance self-play room. A possible interpretation of this pattern of results is that high levels of positive schizotypy produce a bias towards an experience of agency where none is afforded by environmental contingencies; consequently, the zero IT-distance auto-play room is perceived as providing more instrumental control than it actually does, making it more appealing; conversely, the high IT-distance self-play room is rendered less discriminable, in terms of instrumental control, from lower-distance or auto-play rooms, reducing its relative appeal. This interpretation is consistent with work on individuals with schizophrenia, which has shown that positive symptoms of schizophrenia, in particular hallucinations and delusions, are associated with an enhanced sense of agency (12; 13; 16; 14; 15). One possibility is that the preference for greater instrumental divergence may be mediated by a subjective experience of agency, although this characterization is more consistent with the notion of instrumental divergence as a reward surrogate than with the forward planning model found to best predict behavior in the current study.

Measures of the subjective experience of agency vary substantially across studies, from declarative self- vs. external attributions (13; 14; 15; 25), to changes in time perception (12; 19; 26; 27), and interference with motor performance (17). Likewise, ontological accounts of human agency range from philosophical theories of desires and means-end beliefs (28), to socio-cognitive constructs of self-regulation and self-reflectiveness (29). At the core of all these approaches, however, is the notion of an agent as a self-directed cause of internal and external states. Instrumental divergence, as a condition of the environment, is critical to such causation, since it reflects the degree to which an agent can use self-directed actions to generate specific outcome states. The findings reported here suggest that a recently demonstrated preference for environments with greater instrumental divergence, and thus, arguably, greater agency, may reflect an estimation of decision values over a range of possible, dynamic, and potentially aversive, outcome utilities.

Previous work, however, suggests that instrumental divergence is an explicitly represented decision variable: For example, using a simple choice task in which decision outcomes were pictures of various food items to be consumed at the completion of the study, Liljeholm et al. (6) found that BOLD activity in the right supramarginal gyrus was parametrically modulated by changes in instrumental divergence, defined as the IT distance between outcome distributions, across trials. Of course, such neural correlates need not reflect an actual computation of information theoretic variables; Liljeholm (23) noted that the accuracy with which a simple neural network classifies observed outcomes in terms of antecedent actions should scale with the IT distance between outcome distributions associated with action alternatives. Further research is needed to characterize the neural computations mediating an influence of instrumental divergence on goal-directed decision-making.

## Acknowledgements

This work was supported by a CAREER grant from the National Science Foundation (1654187) awarded to Mimi Liljeholm. The author thanks Michelle Liang and Kara Schwartz for assistance with cloud-based data management.

## Supplementary Material

https://osf.io/vqehm/

## Notes

**Conflicts of interest:** There are no conflicts of interests.

https://osf.io/vqehm/

### Competing Interest Statement

The authors have declared no competing interest.

### Summary of Updates

title, text, analyses

